# Design Principles Underlying Robust Adaptation of Complex Biochemical Networks

**DOI:** 10.1101/2020.09.21.307140

**Authors:** Robyn P. Araujo, Lance A. Liotta

**Affiliations:** School of Mathematical Sciences, Queensland University of Technology, Brisbane, QLD, 4000, Australia; Institute of Health and Biomedical Innovation (IHBI), 60 Musk Avenue, Kelvin Grove, Brisbane, QLD, 4059, Australia; Center for Applied Proteomics and Molecular Medicine, George Mason University, 10920 George Mason Circle, Manassas, Virginia, 20110, USA

**Keywords:** Robust Perfect Adaptation, Complexity, Chemical Reaction Networks, Robustness, Network Topology

## Abstract

Biochemical networks are often characterised by tremendous complexity – both in terms of the sheer number of interacting molecules (“nodes”) and in terms of the varied and incompletely understood interactions among these molecules (“interconnections” or “edges”). Strikingly, the vast and intricate networks of interacting proteins that exist within each living cell have the capacity to perform remarkably robustly, and reproducibly, despite significant variations in concentrations of the interacting components from one cell to the next, and despite mutability over time of biochemical parameters. Here we consider the ubiquitously observed and fundamentally important signalling response known as Robust Perfect Adaptation (RPA). We have recently shown that all RPA-capable networks, even the most complex ones, must satisfy an extremely rigid set of design principles, and are modular, being decomposable into just two types of network building-blocks – Opposer modules, and Balancer modules. Here we present an overview of the design principles that characterize all RPA-capable network topologies through a detailed examination of a collection of simple examples. We also introduce a diagrammatic method for studying the potential of a network to exhibit RPA, which may be applied without a detailed knowledge of the complex mathematical principles governing RPA.

## 1. Introduction

The co-existence of both complexity and robustness in the self-organising, self-regulating networks arising in nature represents an extraordinary paradox *[1-3]*. Indeed, since robustness in the face of changing and unpredictable environments is one of the most fundamental requirements for any living system, this raises a deep question about nature’s most basic design principles: How must biological complexity be organised to accommodate the exacting demands of robust performance?

In this chapter we consider this question in the light of the keystone biological function known as Robust Perfect Adaptation (RPA). RPA is the process whereby a system resets selected internal components to their respective pre-stimulus baseline levels (or “set-points”) following a disturbance or altered input, with no need for fine-tuning of system parameters *[4-6]*. The capacity for RPA is widely considered to be an essential characteristic of all evolvable and self-regulating systems *[5]*. It has been ubiquitously observed in biology at all levels of organisation, from intracellular networks comprising genes, metabolites and/or proteins, to signalling networks at the whole-organism level, including calcium or hormone regulation, vision, olfaction, touch, and embryonic morphogenesis *[7-14]*. Dysregulation of RPA is also thought to play a central role in human disease, since maladaptation – the establishing of harmful, disease-promoting RPA set-points – is thought to be a signalling feature underpinning disorders such as drug addiction, chronic pain, cancer progression, and metabolic syndrome (e.g. obesity, high blood pressure and insulin resistance) *[15-20]*.

Early theoretical work in the RPA field focussed on a number of relatively simple RPA-capable network designs, which have attracted widespread attention from the scientific community and have been analysed extensively. Ma et al. *[21]*, for instance, determined that for networks comprising just three interacting nodes, all network arrangements capable of exhibiting RPA to a persistent change in network input can be divided into two well-defined classes – negative feedback loop with buffer node (NFBLB), and incoherent feedforward loop with proportioner node (IFLPN). A highly influential four-node RPA-motif, proposed independently of the work of Ma et al., is the antithetic integral control model *[22,23]*. Recent work has also considered simple RPA-motifs in models that account for dilution of cellular constituents during the cell’s growth phase in order to extend the applicability of existing RPA theory in synthetic biology settings *[24,25]*.

While the study of these small and very specific RPA-promoting network designs has given rise to many important applications of RPA theory in the synthetic biology field *[23,25]*, such simple models are ill equipped to provide insight into the highly complex network designs through which RPA may be realized in nature – that is, in complex *self-organising, self-regulating* and highly mutable systems such as cellular signal transduction networks. It is clear that complex biosystems differ in fundamental ways from engineering control systems, and are comprised of elements that must serve both as the transmitted signals *and their own controllers*. Unlike their designed counterparts in engineering control systems, bionetworks do not have the luxury of employing specially-designed, dedicated components whose purpose is to sense or control biochemical signals. Moreover, these vast and intricate biosystems are typically called upon to solve a large number of “cognitive problems” in parallel – processing and interpreting a high-dimensional space of biochemical and mechanical stimuli, and making decisions as to appropriate systems-level responses. In this context, signalling proteins that play a *regulatory role* (reduplicating a model of the dynamic structure of the input signal) in a *feedback* path, can also simultaneously play a transmissive role (in a “route” leading from input node to output node). Such ambiguous roles for signalling componentry may not be needed, and may not even be appropriate, in an engineering design context.

An understanding of how RPA could be implemented in complex bionetworks amid such strenuous cognitive demands requires access to a “complete” design space for *all possible* RPA-capable network topologies. We recently developed a new mathematical approach to solve this general “RPA problem” *[5]*, and as a consequence, the full set of all possible RPA-capable network configurations is now known – for any sized network, for any degree of complexity (network size and interconnectedness), for any network “type” (e.g. protein network, gene regulatory network, intercellular communication network, neuronal network) and any adaptive time-scale. In particular, we have demonstrated two classes of network topologies – the Opposer Module and the Balancer Module - that can truly be considered topological basis elements, along with a general way to combine those elements, so as to span the complete solution space to the RPA problem.

From this general RPA framework, it has now been established conclusively that RPA-capable networks must be *modular*. The Opposer Module is a rich and potentially complex generalization of the negative feedback integral control that has been well-known to control engineers since the 1970s *[5,26,27]*. The Balancer Module, on the other hand, generalizes the simple incoherent feedforward motif that occurs repeatedly in bacterial transcription networks. It is now clear that the NFBLB motif identified by Ma et al *[21]* is a special case of an Opposer Module, while the IFFLP motif *[21]* is the smallest possible Balancer Module that incorporates an independent “balancer node”. The antithetic integral control motif *[22]* is also a simple Opposer Module. RPA basis modules may be far more complex that these well-studied examples, as we shall see in this chapter, and may be interconnected in well-defined ways to construct arbitrarily large RPA-capable networks *[5]*.

The goal of this chapter is to present the essence of general RPA theory through the discussion and analysis of a selection of examples. Each example is chosen to highlight one or more of the essential design principles that characterise general RPA networks. In each case, we illustrate the use of a diagrammatic method which captures the mathematical content of the “RPA Equation” – the fundamental algebraic condition which must be satisfied by all RPA-capable networks – thus providing a tool for exploring the deep principles of RPA without detailed mathematical knowledge of the underlying theory. It is our hope that this diagrammatic method will offer an accessible overview of RPA theory to mathematicians and non-mathematicians alike, and will stimulate broad scientific interest in this vital topic in biocomplexity theory.

## 2. The Mathematics of RPA

In the interests of a self-contained presentation, we begin with a brief overview of mathematical RPA theory to provide a backdrop for the examples we explore in Section 4. In Section 2.1 we present a brief discussion of some of the earliest known models of RPA networks, which were almost entirely limited to just three or four interacting elements. In Section 2.2, we discuss the topological framework *[5]* that allows more general RPA-capable topologies to be identified, in networks that contain arbitrarily large numbers of interacting elements and interconnections. Readers with limited interest in the mathematical details of RPA in a general network setting may omit this section, and can proceed to the more pragmatic discussion in Sections 3 and 4 without loss of continuity. Section 3 will be devoted to the development of a diagrammatic representation of the “RPA Equation” – the defining algebraic constraint that must be satisfied by all RPA-capable networks. We will employ this diagrammatic method to represent the flow of biochemical information in our example networks later in Section 4, illustrating the essential RPA-promoting mechanisms at play in each case.

### 2.1 Early Models of RPA in Biology

Early approaches to studying RPA may be grouped into three overarching methodologies *[28]*:

1. <u>Development of models corresponding to well-known adaptive systems in biology</u>. One of the best known cases of this strategy is the Barkai-Leibler model of bacterial chemotaxis *[4]*. Barkai and Leibler’s work on chemotaxis in E-Coli has exerted a strong and long-lasting influence on RPA theory for many reasons. It was the first study to show, for a specific molecular signalling network, that RPA could be a systems-level property, independent of parameter fine-tuning. In fact, the use of the word *robustness* to describe the independence of the adaptive performance from system parameters may be traced back to this seminal work. But the signalling events that regulate bacterial chemotaxis, the subject of Barkai and Leibler’s work, are somewhat unusual in that the functionality (in this case, RPA of tumbling frequency) can be captured by only a very small number of signalling molecules. This has reinforced the notion of “motif” in signalling networks – the concept that particular network functionalities such as RPA may be engendered at the level of a relatively small functional unit within a larger network.
2. <u>Ad-hoc modelling</u>. Modelling of small systems of interactions, generally from a nonlinear dynamics perspective *[29]*, has also made significant contributions to early RPA models, often drawing on the possibility of strong parallels between biochemical network structures and engineering design principles. This perspective has produced, among other functional response units, the “sniffer” motif, which is simple two-component RPA model. This modelling framework has also suggested a two-component feedback model for RPA (homeostasis) that relies on an embedded ultrasensitive response element *[29]*, thereby highlighting the deep relationship between ultrasensitivity and RPA. The antithetical integral control motif *[22, 23]* is a more recent example of a small motif structure that produces RPA. This motif has been genetically engineered as a synthetic controller in living cells, and has been applied to growth-rate control in E-coli *[23]*.
3. <u>Computational searches</u>. Comprehensive computational searching strategies consider RPA from the point of view of an inverse problem: Given that the output of the network must return to the same fixed baseline value, irrespective of the input magnitude, what is the space of all possible motifs that could achieve this? This approach is not about asking which solutions are actually observed in particular organisms or signalling contexts, or even about which solutions might be ‘best’; it is purely a matter of delineating the complete space of biochemically plausible ways to implement the required function. The computational demands of this approach *severely* restrict the network sizes that can be investigated by this method, and often impose a level of ‘coarse-graining’ of the parameter space to be sampled. The most comprehensive computational search for solutions to the RPA problem at the present time remains the study by Ma et al. *[21]*, which was limited to small networks of just three nodes. This influential study revealed that for three-node networks, just two types of motif are capable of exhibiting RPA.

Schematic representations of the main classes of small RPA motifs described above are depicted in Figure 1.

**Figure 1:**
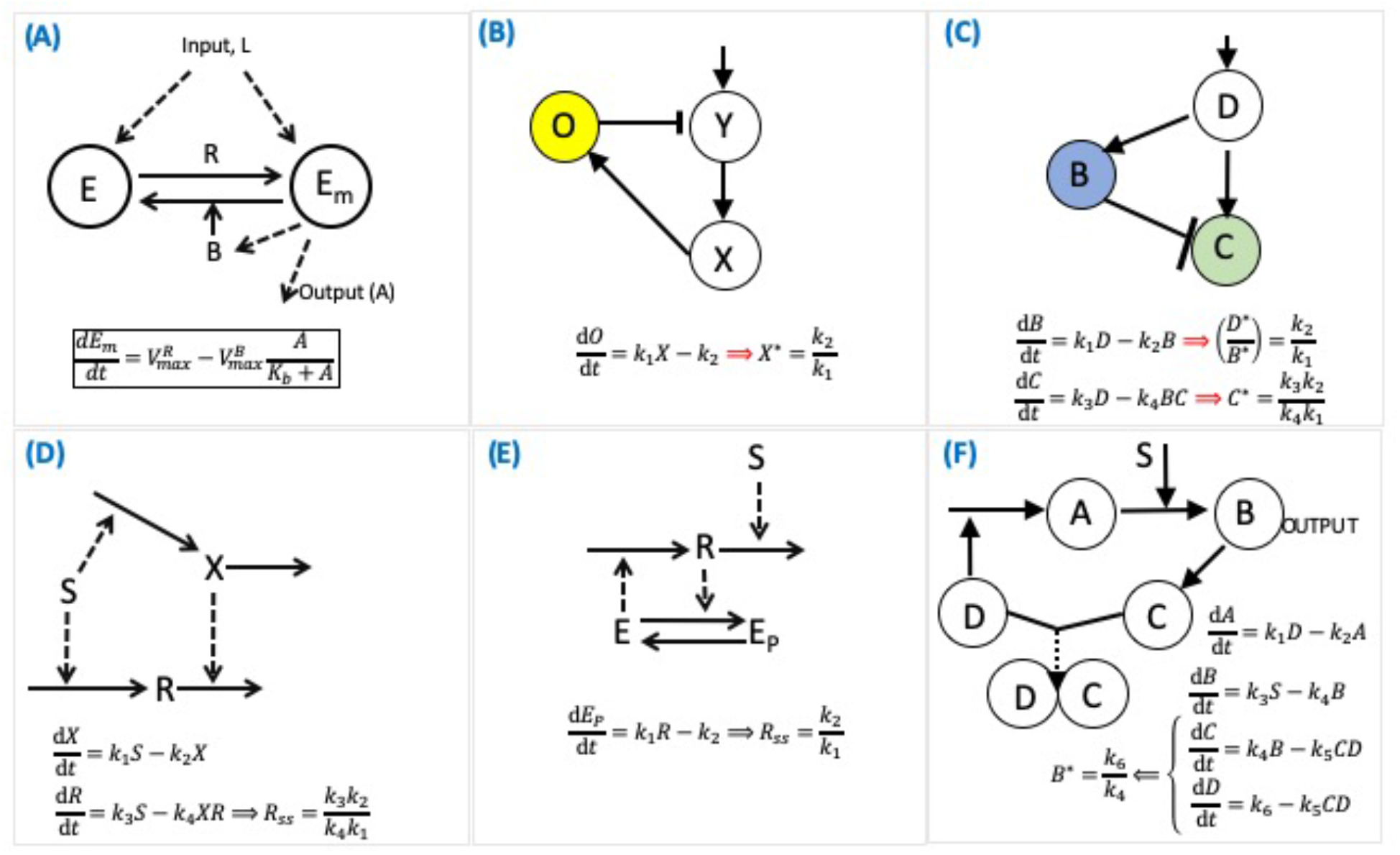
Schematic representations of early RPA models. (A) Barkai-Leibler model of bacterial chemotaxis [4]; (B) Negative feedback loop with buffer node (NFLBN) identified by Ma et al. [21]; (C) Incoherent feedforward loop with proportioner node (IFLPN) identified by Ma et al. [21]; (D) ‘Sniffer’ model of RPA [29]; (E) Homeostasis model with embedded ultrasensitivity [29]; (F) Antithetic integral control model of RPA [22,23].

### 2.2 The mathematical basis for RPA in general interaction networks

We now turn our attention to the more challenging problem of how Robust Perfect Adaptation (RPA) may be achieved in complex networks that are self-organizing, self-regulating and evolvable – in other words, networks that arise in biology. To understand how complexity is organised into robustly adapting systems in nature, it is not a simple matter of looking for *more* solutions to the RPA problem, given that we already know *some* solutions in very small, simple networks (as summarised in Section 2.1). Nor do we seek a method for testing particular network topologies to see if they are capable of RPA. Rather, we seek *the set of* ***all possible*** *RPA network topologies* - that is, all the possible arrangements of nodes that are capable of exhibiting RPA, along with the constraints on reaction mechanisms at those nodes.

Our recent work *[5]* has demonstrated that two – and only two - classes of network topology (modules) can truly be considered *topological basis elements* for RPA in arbitrarily complex networks. These topological basis modules, along with a general way to combine those modules, *span* the complete solution space to the RPA problem. RPA-capable networks are thereby constrained to be modular. This fundamental property of robustly performing complex networks suggests that they may now be studied from the point of view of their unexpected simplicity – that is, as decompositions into basis modules. Since the set of all possible network arrangements grows factorially with network size, the very limited set of RPA-capable network topologies defined by this basis effectively represents the “needles in the haystack” that evolution must seek out over and over again as networks that require the RPA property grow and change over time.

The essential foundation for the development of this more RPA theory may be summarised briefly as follows (see *[5]* for full details):

Suppose we have *n* interacting ‘nodes’. Typically, ‘nodes’ refer to the interacting physical components in the system although, they could refer to more complicated mathematical expressions involving these interactions *[5]*. Here, for the sake of concreteness, it will be helpful to consider the ‘nodes’ to be the interacting proteins in an intracellular signalling network. We denote the respective concentrations (or abundances) of these interacting proteins as *P*_*1*_, *P*_*2*_, …, *P*_*n*_. One of these proteins, the ‘input node’, will receive a signal from outside the network, while one node (often, but not necessarily, distinct from the input node) is the endpoint of interest, which is expected to exhibit RPA in response to the signal received at the input node, and is given the status of the ‘output node’. Let us assume that the *n* nodes are ordered so that the input node is *P*_*1*_ and the output node is *P*_*n*_. For a network with single input/output node, we will place this node first in the ordering, *P*_*1*_.

Now, to each node *P*_*i*_ in the set, we associate a reaction rate *f*_*i*_ *= dP*_*i*_ */dt*, which allows us to account for the nature and strength of the interactions among the *n* nodes of the network. The rate *f*_*i*_ will always be regulated by at least one *other* node, say *P*_*1*_. It will almost always also depend on the node’s own concentration, *P*_*i*_, except in the case of an exceptional reaction mechanism of central importance to the RPA problem – the “opposition mechanism” – which we discuss in Section 3.2. The input node, *P*_*1*_, will also be regulated by the external stimulus, *S*. In general, *f*_*i*_ could depend on any subset of the n network nodes. Thus, *f*_*i*_ has the general form

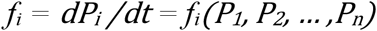

for all nodes not receiving an external stimulus, and

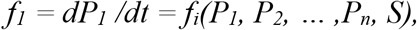

for the input node, *P*_*1*_.

It is a straightforward algebraic problem to determine that there are two key mathematical quantities that codify the sensitivity of the output node’s steady-state to the level of the externally-delivered stimulus. These are (1) the system Jacobian, *J*_*n*_, and (2) the “input-output minor”, *M*_*1n*_, associated to *J*_*n*_. We depict the structures of these quantities in Figure 2. As shown, the input-output minor *M*_*ij*_ is obtained from *J*_*n*_ by eliminating the row associated with the network’s input node, *P*_*i*_, and the column associated with the output node, *P*_*j*_.

**Figure 2:**
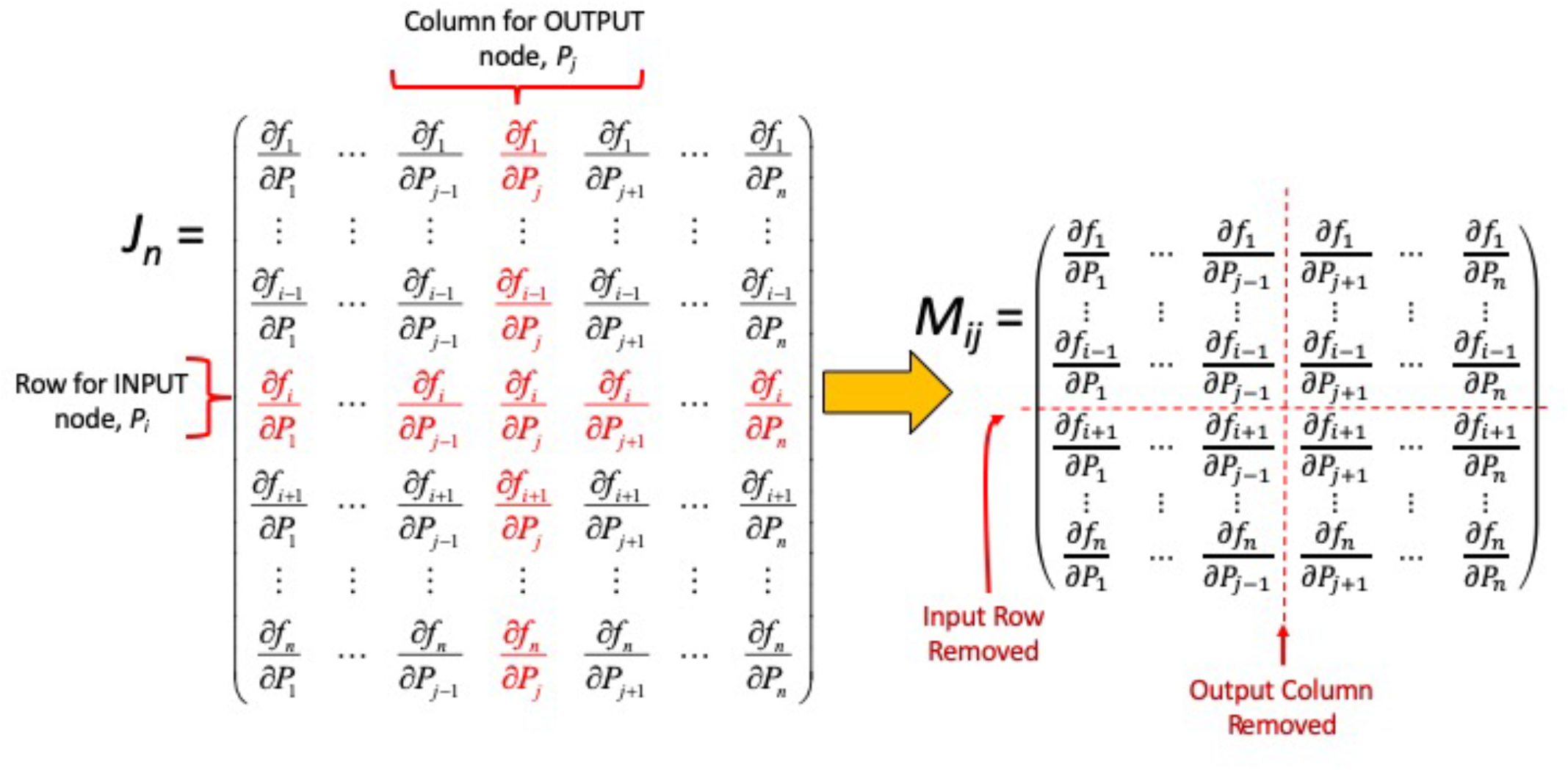
The RPA Equation – the fundamental algebraic condition that must be satisfied by all RPA-capable networks – is derived from the ‘input-output’ minor, M_ij_, associated to the input node (P_i_) and output node (P_j_) for a network. As indicated, M_ij_ is obtained from the Jacobian matrix, J_n_, of the system by eliminating the row of J_n_ associated with the input node, P_i_, and the column associated with the output node, P_j_.

The ratio of the determinants of these two key matrices gives rise to the special algebraic condition which must be satisfied by all RPA-capable networks *at steady-state*, for all possible parameter sets, and all possible stimulus levels delivered to the system. In particular,

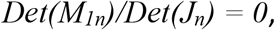

Or, equivalently,

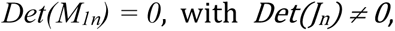

is a necessary condition for RPA in all networks, regardless of size or complexity, and must be satisfied by all system steady states, for all possible parameter choices, and all stimulus levels. We refer to the equation *Det(M*_*1n*_*)=0* hereafter as ***the RPA equation***, with the condition *Det(J*_*n*_*) ≠ 0* denoted ***the RPA constraint***. Rigorous analysis of these algebraic conditions *[5]* has shown that the RPA equation, in particular, induces tremendous structure on network designs that accommodate the RPA property.

Note that both these mathematical conditions can contain an extraordinarily large number of terms in networks containing more than just a handful of nodes. Indeed, the RPA equation can contain up to *(n-1)!* terms, depending on how extensively the *n* nodes are interconnected. The RPA constraint can contain up to *n!* terms. Fortunately, a topological approach to the structure of the RPA equation *[5]* has simplified this problem tremendously, and allowed the set of all possible RPA-capable network configurations to be determined – not by enumeration, of course, but through the identification of a suitable basis.

With a view to describing the essential nature of this (topological) basis, along with the rules for combining (interconnecting) basis elements in general RPA-capable networks, we devote the next section to a new diagrammatic method for presenting the mathematical content of the RPA equation.

## 3. Methods for RPA Analysis

### 3.1 A diagrammatic method for analysing the RPA capacity of a network

As is evident from the discussion in the previous section, we require remarkably little biochemical information about a particular network to analyse its capacity for RPA. The RPA equation comprises a summation of terms, each a product of *(n-1)* factors of the form ∂*f*_*i*_/∂*P*_*j*_ or the form ∂*f*_*k*_/∂*P*_*k*_. Each factor of the form ∂*f*_*i*_/∂*P*_*j*_ corresponds to a network interconnection *P*_*j*_ →*P*_*i*_, while ∂*f*_*k*_/∂*P*_*k*_ may be referred to as the ‘kinetic multiplier’ *[5]* associated to the node *P*_*k*_. As we will see, kinetic multipliers play a pivotal role in ‘solving’ the RPA equation since these are the only class of factors that can be identically zero while still preserving the configuration of the underlying network. In particular, a zero value for a factor of this type places constraints on the reaction mechanism occurring at the node in question, without removing any network linkages impinging on the node.

The mathematical content of each term in the RPA equation thus suggests a diagrammatic representation: Replacing the factor ∂*f*_*i*_/∂*P*_*j*_ with the symbol *P*_*j*_ →*P*_*i*_, and representing ∂*f*_*k*_/∂*P*_*k*_ with the symbol (*P*_*k*_), we see that the terms of the RPA equation correspond to successions of linkages, along with kinetic multipliers, that exist in the underlying network. Together, these form ‘routes’ leading from the input node to the output node, feedback loops (multi-node cycles), and products of kinetic multipliers (one-node cycles).

In fact, we have shown *[5]* that the special algebraic structure of the determinant map will constrain each term of the RPA equation to assume the following form:

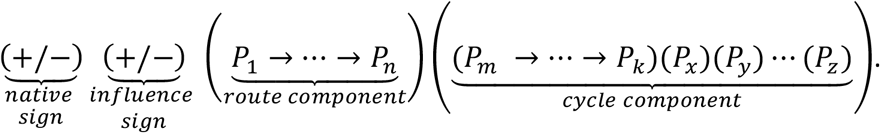

In the special case of a network with a single input/output node, the route component will reduce to unity, producing an RPA equation with only cycle components:

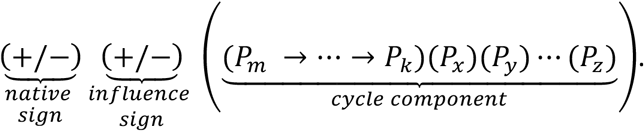

In fact, each and every ‘route’ in a particular network (leading from the input node through to the output node) will be represented as a route component *P*_*1*_ *→ … → P*_*n*_ in the associated RPA equation. Likewise, each and every feedback loop that exists in the network will appear in the cycle component of the RPA equation. In particular, the full set of cycle components appended to a particular route component will be generated by *all possible combinations of cycles* (i.e. feedback loops and kinetic multipliers) formed by the nodes that are *disjoint from* (not occurring in) that route component. For clarity and convenience, we represent a feedback loop of the form *P*_*a*_ *→ P*_*b*_ *→ P*_*c*_ *→ P*_*a*_ with the diagrammatic representation (*P*_*a*_ *→ P*_*b*_ *→ P*_*c*_) – see Figure 3. The RPA equation may thus be interpreted as a summation of all possible route-cycle combinations for its network.

**Figure 3:**
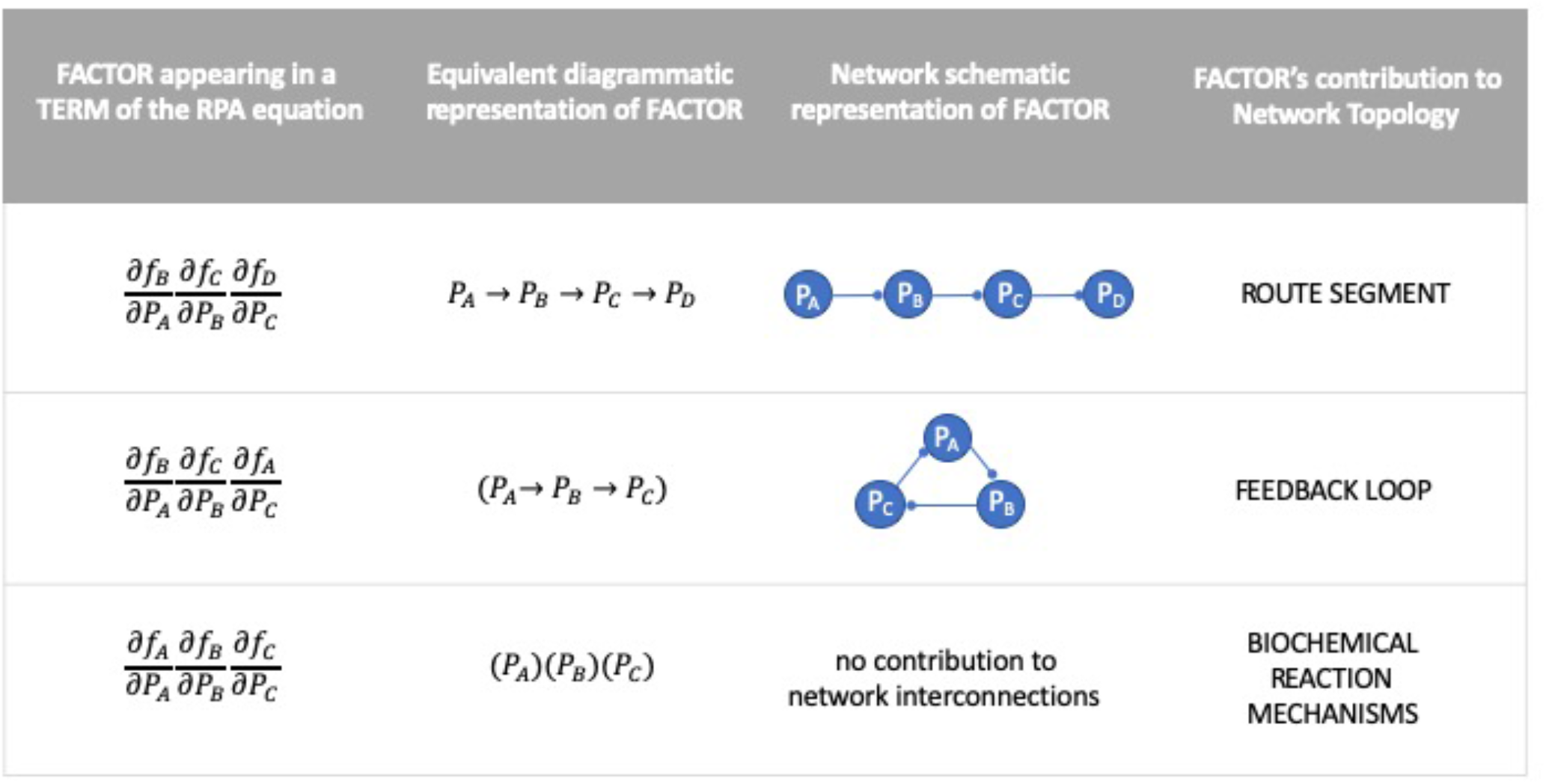
Collections of factors appearing in a single term of the RPA equation may be represented diagrammatically as route segments, feedback loops or one-node cycles (“kinetic multipliers”). Of these three types of network contributions resident in the terms of the RPA equation, only kinetic multipliers can assume a zero value without affecting the interconnectivity of nodes in the network, since these reflect the nature of the reaction mechanisms at individual nodes.

Note that each term in the RPA equation is prefixed with a positive or negative sign. There are two contributing factors to this sign, so we represent it as the product of two component signs – the *native sign* and the *influence sign*, as indicated above. The native sign is inherited from the term’s contribution to the determinant expansion. It is straightforward to show (see Proposition 1 in *[5]*) that the native sign for the term is given by (−1)^*z*+*n*+1^, where *z* is the number of distinct cycles (feedback loops plus kinetic multipliers) present in the term, and *n* is the number of nodes in the network. The influence sign, on the other hand, takes account of how many inhibitory interactions occur in the sequences of node-node linkages comprising routes and feedback loops represented in the term. An odd number of inhibitory interactions attracts a negative sign, while a positive number of such interactions attracts a positive sign. Note that kinetic multipliers always attract a negative sign *[5]*.

In Figure 4 we demonstrate the application of these simple principles by deducing the diagrammatic representation of the RPA equation for a simple 6-node network. For reasons that will become clear through the analysis of the examples in Section 4, the particular network configuration depicted in Figure 4 is *not* capable of exhibiting RPA. Note that here, and in all subsequent network schematics, we adopt the convention that the symbol → represents an *activating* interaction while the symbol —• denotes an *inhibitory* interaction. For this network, *P*_*1*_ is selected as the input node, while *P*_*6*_ is the output node.

**Figure 4:**
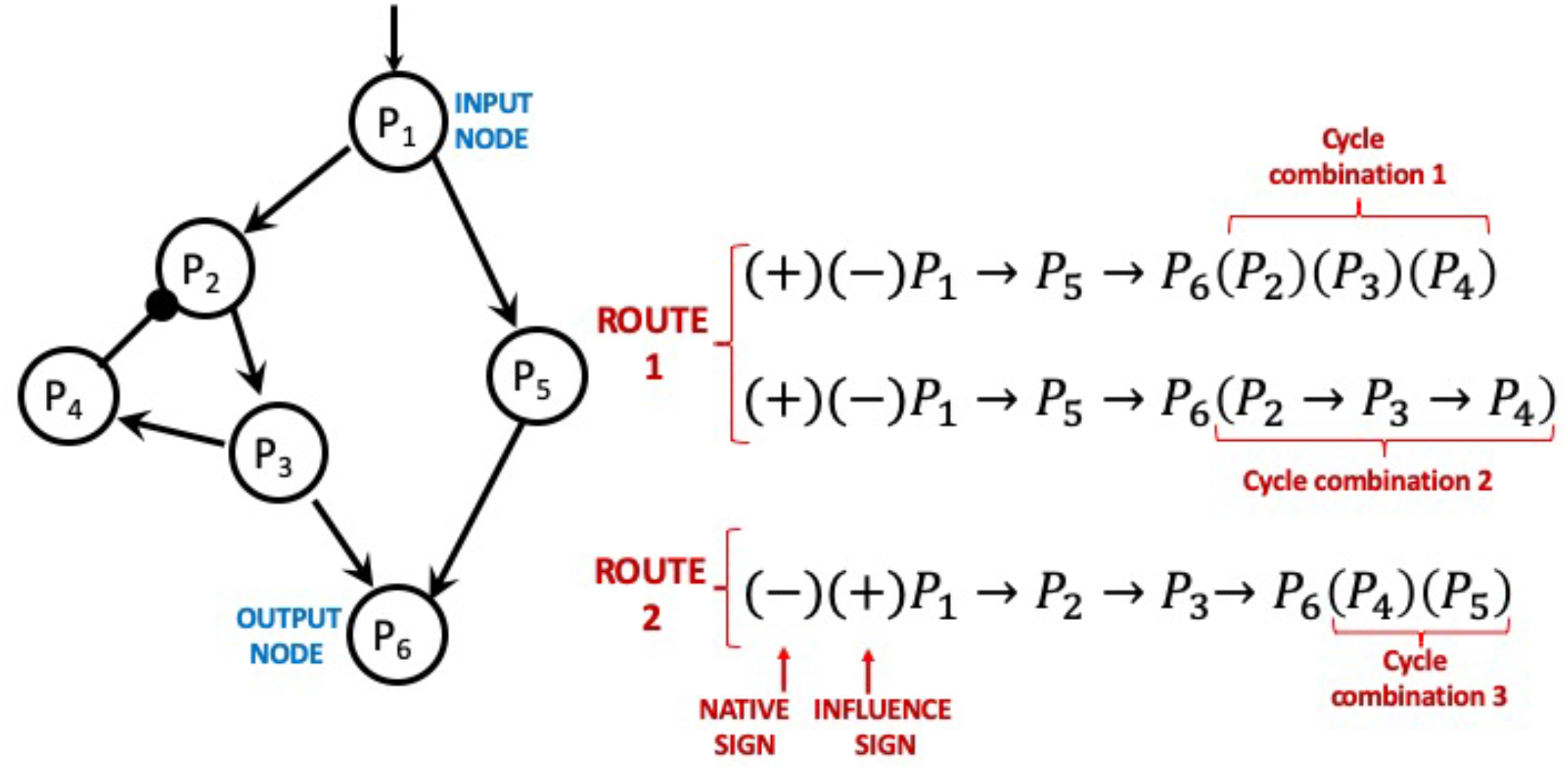
A simple six-node network example, illustrating the diagrammatic method for deducing the RPA equation for a network. As shown, this network comprises two distinct routes from input (P_1_) to output (P_6_). The cycle combination appended to each route comprises all possible combinations of cycles composed of nodes that are disjoint from that route.

In this particular network, there are two distinct routes from *P*_*1*_ to *P*_*6*_: *P*_*1*_*→P*_*5*_*→P*_*6*_, and *P*_*1*_*→P*_*2*_*→P*_*3*_*→P*_*6*_. For the route *P*_*1*_*→P*_*5*_*→P*_*6*_, containing no inhibitory interactions, there is just one feedback loop disjoint from the route, namely (*P*_*2*_*→P*_*3*_*→P*_*4*_). There are thus two possible cycle combinations appended to this route: (*P*_*2*_*→P*_*3*_*→P*_*4*_) and (*P*_*2*_)(*P*_*3*_)(*P*_*4*_). Both combinations contribute a negative interaction sign to the term (a *negative* feedback loop, and an *odd* number of kinetic multipliers). The native sign contributed by the first combination is (−1)^1+6+1^ = (+1); for the second combination, the native sign is (−1)^3+6+1^ = (+1).

For the route *P*_*1*_*→P*_*2*_*→P*_*3*_*→P*_*6*_, also containing no inhibitory interactions, there are no disjoint feedback loops; the route is therefore only appended to the cycle combination (*P*_*4*_)(*P*_*5*_). This combination contributes a positive interaction sign (an *even* number of kinetic multipliers), with native sign equal to (−1)^2+6+1^= (−1).

### 3.2 Fundamental design principles of RPA networks

Equipped with an understanding of the mathematical content of the RPA equation, we may now consider the general properties of network configurations that satisfy the RPA equation. At the broadest level, identifying the full space of all RPA-capable network topologies relies on a general method for partitioning the RPA equation to ‘independently-adapting subsets’. This framework interprets the terms of the RPA equation as elements of a topological space, with the independently-adapting sets serving as the closed sets of that space *[5]*. Such partitions of the RPA equation impose severe constraints on the design principles of networks capable of exhibiting RPA.

With the aim of presenting a non-technical exposition of the essential design principles of RPA-capable networks uncovered by this approach, we proceed in two steps: First, we present (below) a summary of the network design principles that may be deduced from the properties of a mathematically-valid partition of the RPA equation. Then, in the remainder of this chapter (Section 4), we illustrate the application of these core RPA design principles through the discussion and analysis of five simple network examples, exploiting the use of the diagrammatic method we presented in Section 3.1 The design summary below and the set of illustrative examples together give a comprehensive overview of the essential design features that characterise all RPA-capable networks.

#### SUMMARY OF DESIGN PRINCIPLES OF RPA-CAPABLE NETWORKS

A. *General Principles*
  a. There exist only two, and only two, distinct types of independently-adapting subsets in a mathematically valid partition of the RPA equation: S-sets and M-sets. All terms contained in an S-set are identically zero due to the presence of a “special” kinetic multiplier (*P*_*k*_) ≡ ∂*f*_*k*_/∂*P*_*k*_ = 0 (at steady-state) for all stimulus levels, and for all network parameters. All terms contained in an M-set are strictly non-zero, and exactly ‘balance’ – for all stimulus levels, and for all network parameters.
  b. In a valid partition, terms are assigned to independently-adapting subsets *by route*. That is, once a term has been assigned to a particular set in the partition, all terms with the same route component must also been assigned to that set.
  c. The terms of an S-set correspond to the presence of an Opposer Module in the associated network. The special reaction mechanism (*P*_*k*_) ≡ 0 referred to as an Opposer mechanism, or Opposer kinetics. The node *P*_*k*_ is referred to as an Opposer node. Opposer nodes can *only* exist within a feedback loop (see Theorem 1 in *[5]*). The essential structure of an Opposer Module is illustrated in Figure 5. The feedback architecture is defined by a diverter node (D) at the base of the module, and a connector node (C) at the apex of the module.
  d. The terms of an M-set correspond to the presence of a Balancer Module in the associated network. As illustrated in Figure 6, a Balancer Module is defined by a diverter node (D) at the apex, and a connector node (C) at the base, with one or more route segments connecting these two nodes. Subnetworks of arbitrary complexity may be embedded into these route segments. All nodes downstream of the diverter node, and upstream of the connector node, are referred to as Balancer Nodes.
  e. A network for which the input and output nodes are *not distinct* (i.e. a network with a single input/output node) has an RPA equation which may *only* contains S-set(s). Equivalently, a network with a single input/output node can *only* achieve RPA through a decomposition into one or more Opposer Modules. It cannot contain Balancer Modules (see Example Network 3 in Section 4).
B. *Opposer Principles*
  a. An Opposer Module can function with a single Opposer node, but can also employ a collection of Opposer nodes which work collaboratively to ‘solve’ the RPA problem for the network. Such a collection of opposer nodes is known as an ‘opposing set’. Opposing sets must work within a very well-defined sub-network topology, embedded into a feedback loop (see Theorem 3 in *[5]*). Some simple configurations for valid opposing sets are depicted in Figure 5. Interested readers are referred to *[5]* for a more extensive description of opposing set topologies.
  b. An Opposer node requires a *unique* regulator node. If, in practice, multiple nodes regulate an Opposer Node, then this unique regulator is the nearest upstream branchpoint of these multiple regulatory pathways (see *[5]*). For clarity, Section 4 only considers examples where the Opposer Node is regulated directly by a single upstream node.
  c. An Opposer node exhibits Opposer kinetics (see point *c* under ‘General Principles’ in part A above) via a mathematical principle known as *kinetic pairing* (see [5] for further details). If *P*_*o*_ is the concentration of the opposer node, and *P*_*R*_ is the concentration of its unique regulator, then kinetic pairing results in the vanishing of a function of the following form on the steady-state locus of the system: g(*P*_*o*_).(*P*_*R*_ – *k*), where *k* is a function of network parameters. The simplest possible function form for Opposer kinetics is a rate equation that is zero-order in *P*_*o*_, which is the functional form we adopt in all our examples in Section 4. Much more complicated mechanisms for orchestrating Opposer kinetics are possible however (see *[5]* for a more extensive discussion of this point).
  d. An Opposer node never exhibits the RPA property. Rather, an Opposer Node uses kinetic pairing (see point *c* above) to compute an ‘error integral’ *[5]* which confers the RPA property on its unique regulator node. This, in turn, confers the RPA property on various other nodes within the Opposer Module (see Figure 5).
  e. An Opposer node can never directly regulate another Opposer node. This fact places severe constraints on the topologies of opposing sets (see point *a* above).
C. Balancer Principles
  a. In order to generate a Balancer Module, each M-set requires at least one term with negative sign, and at least one term with positive sign.
  b. Balancer nodes and Connector nodes are subject to special reaction mechanisms – the Balancer mechanism and the Connector mechanism, respectively. If *P*_*B*_ is the concentration of a Balancer node, and *P*_*RB*_ is the concentration of its unique regulator, then kinetic pairing results in the vanishing of a function of the following form on the steady-state locus of the system: (*P*_*B*_/*P*_*RB*_ – *k*_*B*_), where *k* is a function of network parameters. If *P*_*C*_ is the concentration of a Connector node, and *P*_*D*_ is the concentration of the Diverter node of the module, then kinetic pairing results in the vanishing of a function of the following form on the steady-state locus of the system: g(*P*_*D*_).(*P*_*C*_ – *k*_*C*_), where *k*_*C*_ is a function of network parameters.
  c. Balancer nodes exhibit Balancer kinetics, and Connector nodes exhibit Connector kinetics, via the mathematical principle known as kinetic pairing (see point c under ‘Opposer Principles’ in part B above).
  d. The presence of the requisite term signs (see point *a* above) and the requisite reaction mechanisms at balancer/connector nodes enables the M-set to achieve independent adaptation through a ‘balancing act’ *[5]*.
D. Principles of Intermodular Connections
  a. AS noted schematically in Figures 5, 6 and 7, some nodes with an RPA module exhibit RPA, and some do not. Connector nodes and Opposer regulators are examples of nodes that do exhibit RPA, while Balancer nodes and Opposer nodes are examples of nodes that do not. Outgoing regulations from nodes that exhibit RPA are referred to as ‘blind’ regulations. Any new network linkages that emanate from these nodes, leading ultimately to the output node, merely supply additional terms to the existing subset of the RPA equation partition associated with the module in question. The new network connections are thus to be interpreted as part of the exist module. By contrast, outgoing regulations from nodes that do not exhibit RPA are referred to as ‘live’ regulations. Any new network linkages that emanate from live nodes supply new terms that must be assigned to a different subset in the partition of the RPA equation. These new ‘routes’ must therefore be connected downstream to a separate RPA module. The two modules act independently, and are said to be connected “in parallel” (see Network Example 1 in Section 4).
  b. It is possible for a single subset in a partition of the RPA equation to be associated with multiple potential modular structures in the underlying network. In this case, the multiple modules are said to be connected “in series” (see Network Example 2 in Section 4).

**Figure 5:**
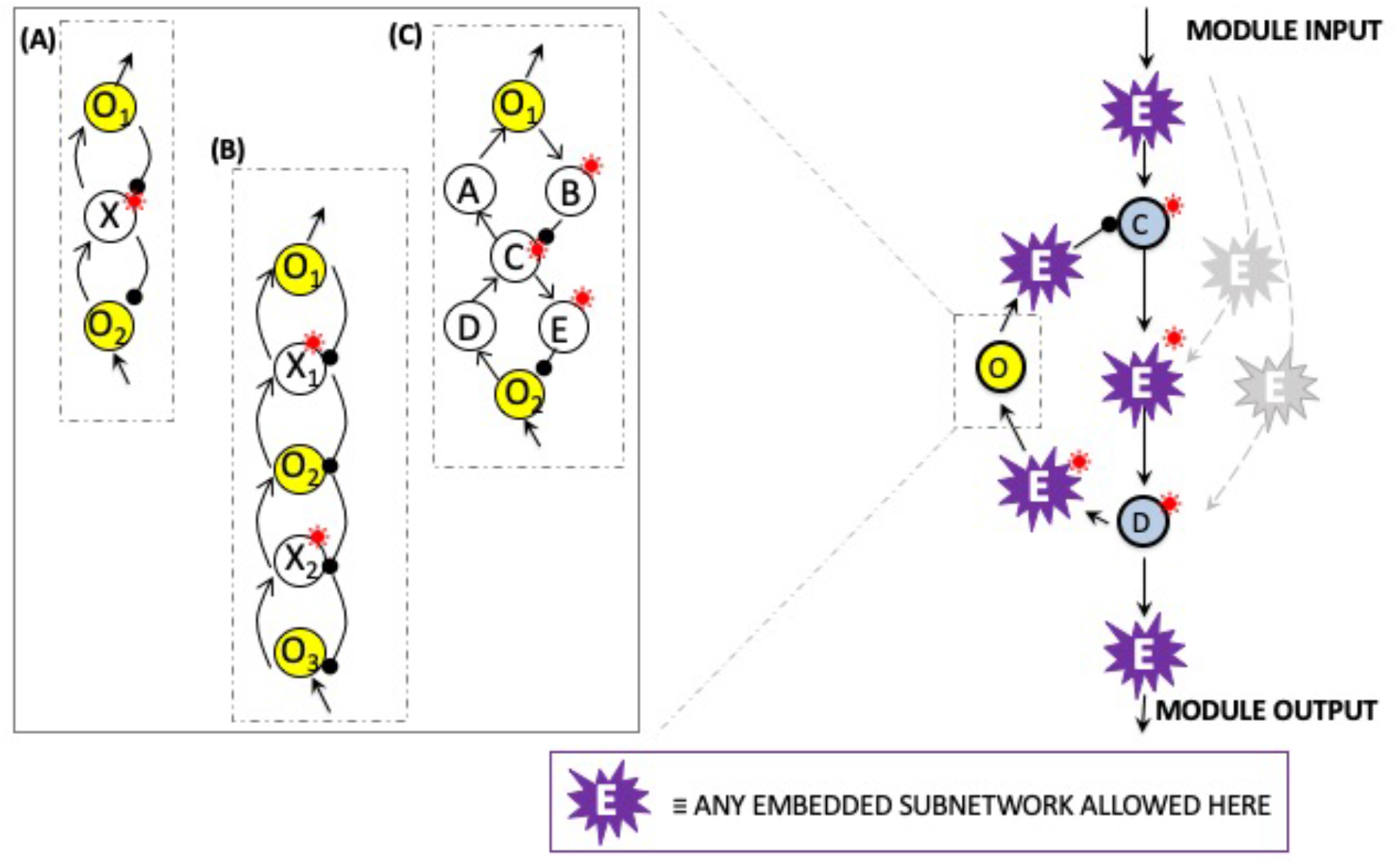
Essential design features of an Opposer module. Opposer modules are characterised by a feedback architecture delineated by two nodes – a connector node, C, at the apex of the module, and a diverter node, D, at the base of the module. Arbitrary subnetworks may be embedded at the positions indicated without affecting the RPA-promoting function of the module. At least one computational node (an Opposer node) is embedded into the feedback segment of the module. All nodes that exhibit the RPA property due to the activities of the Opposer node(s) are indicated with a red asterisk. Collections of opposer nodes may be organised in intricate topological arrangements within the feedback segments of Opposer modules, and work together collaboratively to confer RPA on the diverter node of the module. Several possible opposer node arrangements are illustrated in the inset to the figure (see [5] for a more complete description of ‘opposing sets’ and their associated ‘master sets’): (A) a two-node opposing set with three-node master set; (B) a three-node opposing set with five-node master set; (C) a two-node opposing set with seven-node master set.

**Figure 6:**
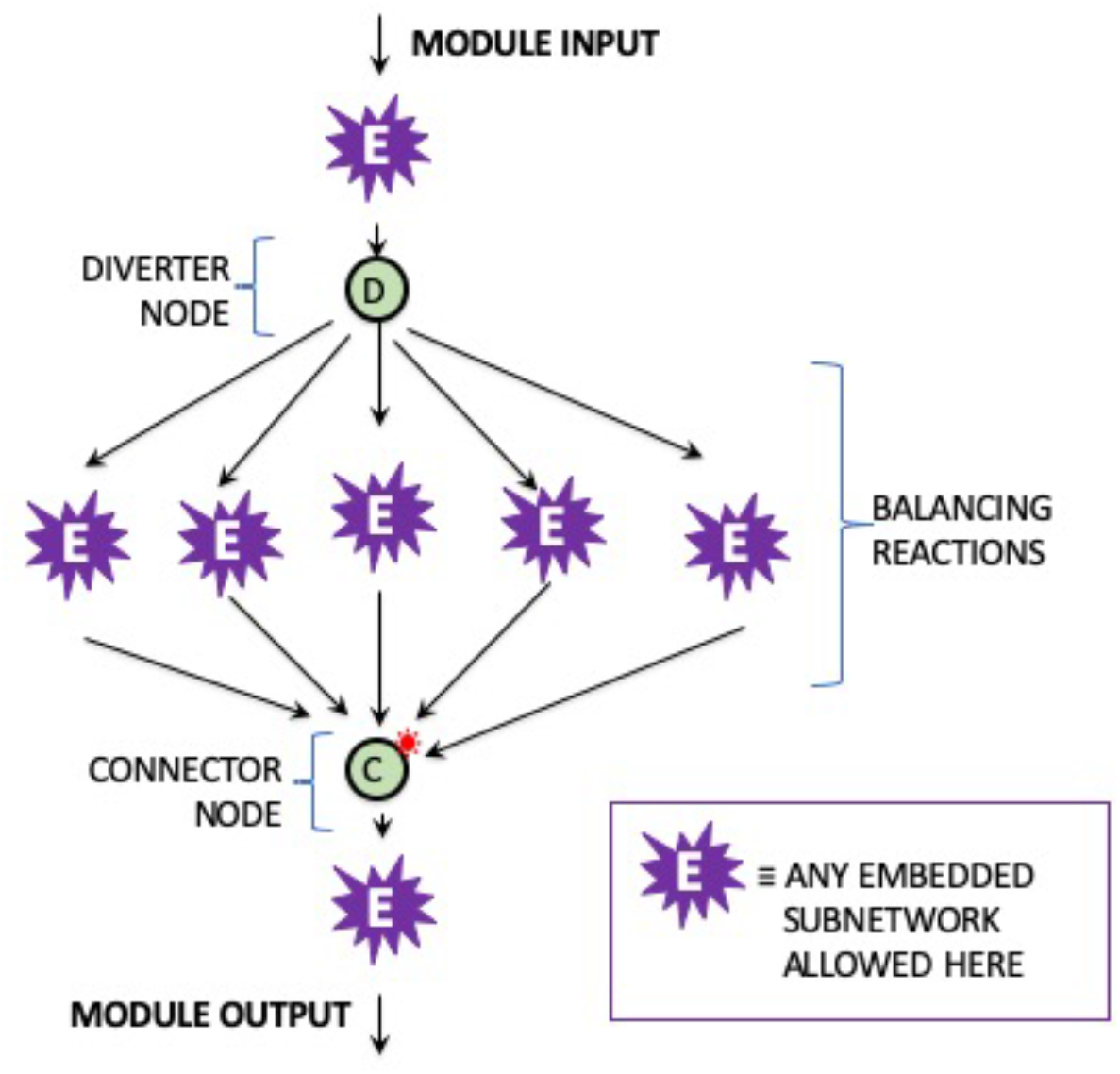
Essential design features of a Balancer module. Balancer modules possess a ‘feedforward’ topology, which may incorporate any number of parallel pathways between the two defining nodes of the module – the diverter node, D, at the apex, and the connector node, C, at the base. Feedback loops may also be embedded into these parallel pathways, as indicated. Only the connector node exhibits the RPA property in a Balancer module, as indicated by the red asterisk.

A final comment on stability is in order before proceeding to our network examples. Identifying RPA-exhibiting steady states from the solution of the RPA equation is not, in itself, a guarantee that the associated ‘set-point’ is a stable one. System stability is, in general, a complex matter that is beyond the scope of the present chapter, and for which we provide some guidance in *[5]*. We do note here that in the context of feedback regulation, positive feedback is highly destabilizing while negative feedback, although no guarantee of stability, is stability-promoting. For this reason, all feedback loops chosen in the following examples are negative feedback loops, for which we show that stable RPA performance readily obtains.

**Figure 7:**
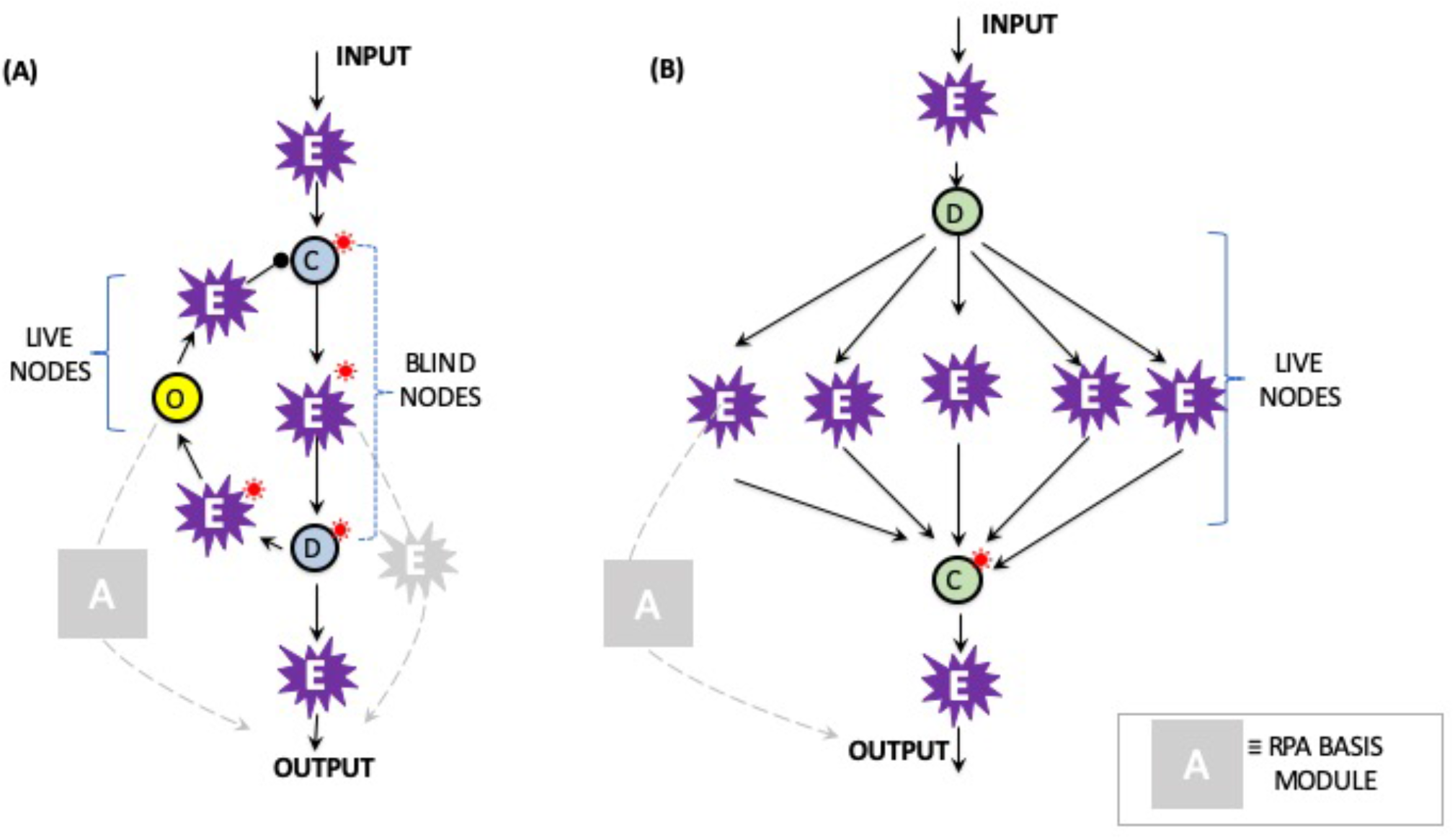
In the context of intermodular connectivities, nodes that exhibit the RPA property within an RPA module are referred to as ‘blind nodes’, while nodes that do not exhibit the RPA property are known as ‘live nodes’. Any live nodes that contribute to a route of the network must be connected downstream to another RPA module (as indicated schematically by the symbol A above). Blind nodes can contribute to additional routes without affecting the RPA capacity of the network.

## 4. Applications of the Diagrammatic Method to Example Networks

In this Section we illustrate these topological principles through an examination of five simple networks, each having an RPA equation containing only a small number of terms. In each case, we examine the conditions under which the network could exhibit RPA, identifying the computational nodes that are subject to special constraints on reaction mechanism in order to solve the RPA equation.

### 4.1 Example Network 1: An opposer module in parallel with a balancer module

In Figure 8 we depict a simple five-node network that receives an external stimulus at P_1_. Nominating the node P_5_ as the output, this network contains a number of parallel pathways, as well as a negative feedback loop between nodes P_1_ and P_2_. Appealing to the diagrammatic method introduced earlier, we readily conclude that the RPA equation for this particular network configuration contains three terms, as shown in Figure 8.

**Figure 8:**
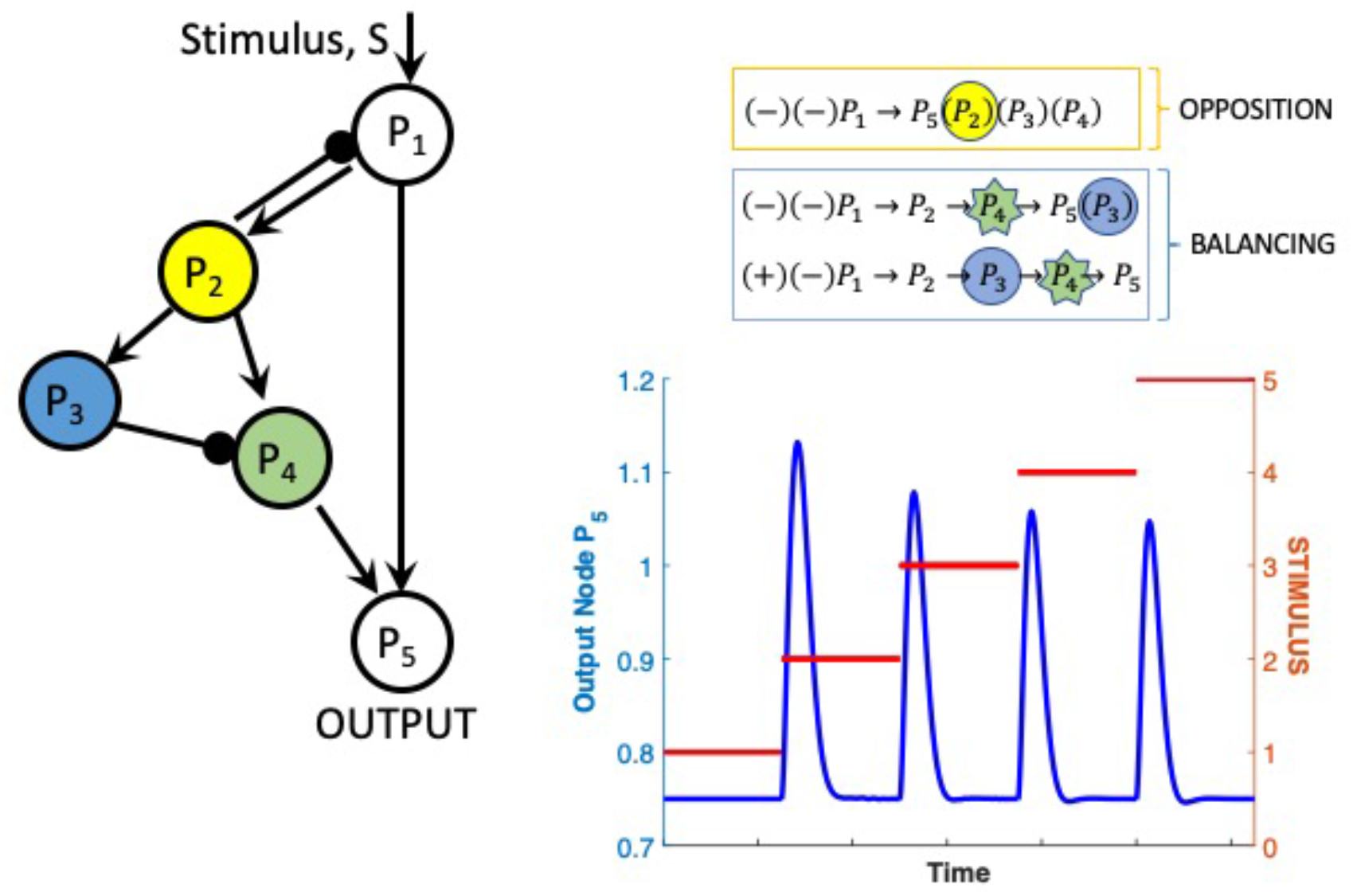
Network Example 1 – a two-node Opposer Module in parallel with a three-node Balancer Module. Parameters: k_1_ = k_3_ = k_4_ = k_10_ = 1; k_2_ = k_5_ = k_6_ = k_7_ = k_8_ = k_9_ = k_11_ = k_12_ = 0.5.

A simple partition of the RPA equation for this network may achieved as follows: A two-node opposer module is formed through the feedback interactions between P_1_ and P_2_. The node P_2_ plays the role of the opposer node (noted in yellow), and is subject to Opposer kinetics 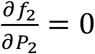. The single regulator of this opposer node, P_1_, will thus exhibit RPA under these conditions, and the first term of the RPA equation noted in Figure 8 will be assigned to an S-set (noted in yellow).

But the Opposer node, P_1_, also appears in a *route* of the network: The network connections P_2_→P_3_ and P_2_→P_4_ are therefore ‘live’ regulations, and the terms containing these regulations in their route components must be assigned to a separate subset in the partition. We see that two different pathways emanating from P_2_ reconnect at node P_5_, and noting that the two remaining terms of the RPA equation have different signs, it follows that assigning P_3_ the role of a Balancer node (noted in blue), and P_4_ the role of a Connector node (noted in green), will allow P_4_ to exhibit RPA. These terms are thereby assigned to an M-set (noted in blue in the diagrammatic representation). Under these conditions, the output node, P_5_, is regulated by two RPA-exhibiting nodes (P_1_ and P_4_), and must, itself, exhibit RPA.

Thus, for this RPA solution, there are three nodes subject to special reaction mechanisms: P_2_ requires Opposer kinetics, P_3_ requires Balancer kinetics, and P_4_ requires Connector kinetics. The remaining nodes (P_2_ and P_1_) are not subject to any constraints; the RPA-capacity of this network is unaffected by the reaction kinetics at these nodes. For illustrative purposes, we choose the simplest possible functional forms for the rate equations at P_2_, P_3_ and P_4_ as follows:

For Opposer node, P_2_:

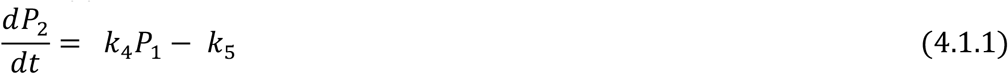

Since this reaction rate is zero-order in P_2_, it is clear that the Opposition condition 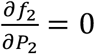 will obtain at steady state, for any choice of parameters. In fact, it readily follows from this equation that the unique regulator node, *P*_1_, exhibits the RPA property with set-point *k*_5_/*k*_4_.

For Balancer node, P_3_:

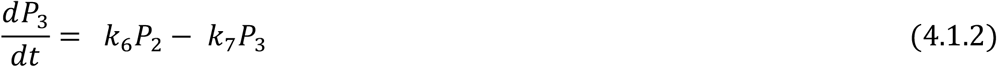

This rate equation ensures that P_3_ will always be proportional to P_2_ at steady state:

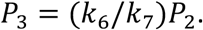

For Connector node, P_4_:

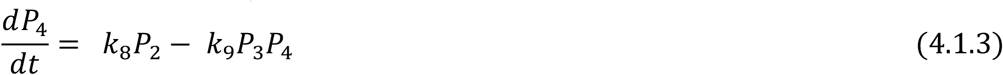

Connector kinetics are ensured by this equation, since both terms are ‘balanced’ by the diverter node, since at steady-state, *P*_3_ = (*k*_6_/*k*_7_)*P*_2_ due to the upstream balancer kinetics. It follows that P_4_ exhibits the RPA property with set-point *k*_8_*k*_7_/*k*_9_*k*_6_.

As noted above, any functional forms are allowed for the remaining nodes, provided the reaction rates accurately reflect the regulations noted in the network schematic. We choose the following simple forms:

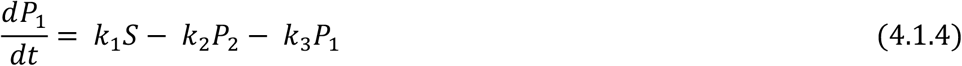

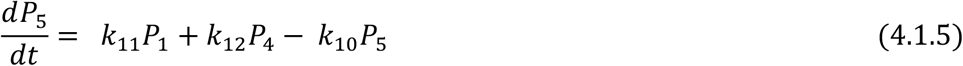

This network thus achieves RPA through the actions of two independently functioning modules – an Opposer module, and a Balancer module. Since the modules contribute independently, each associated to its own subset in the partition of the RPA equation, the modules are said to be *in parallel* (see [5] for further discussion).

We solve Equations (4.1.1) through (4.1.5) above for a range of different stimulus strengths, with results as shown in Figure 8 (see Figure caption for parameter values). Notice that each time the stimulus (red curve) undergoes a step increase, the output node (blue curve) exhibits a rapid initial increase, but then diminishes back to the baseline level, even though the larger stimulus magnitude persists. Since the output always returns to the same value, regardless of the stimulus level being received at the input node, the system exhibits RPA.

### 4.2 Example Network 2: An opposer module in series with a balancer module

Figure 9 presents an alternative combination of the two module types considered in the previous example. In this configuration, the two-module structure is, in fact, *redundant*. In particular, this network will exhibit RPA as a single opposer module if the node P_2_ acts as an opposer node (i.e. if it operates with opposer kinetics 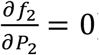). If P_2_ does not act as an opposer, the network can still exhibit RPA as a single balancer module, provided P_3_ exhibits balancer kinetics, and P_4_ exhibits connector kinetics. In other words, this particular configuration can operate *either* as an Opposer module *or* as a Balancer module. For this reason, we say that the two potential modules – the Opposer and the Balancer – are topologically *in series*.

**Figure 9:**
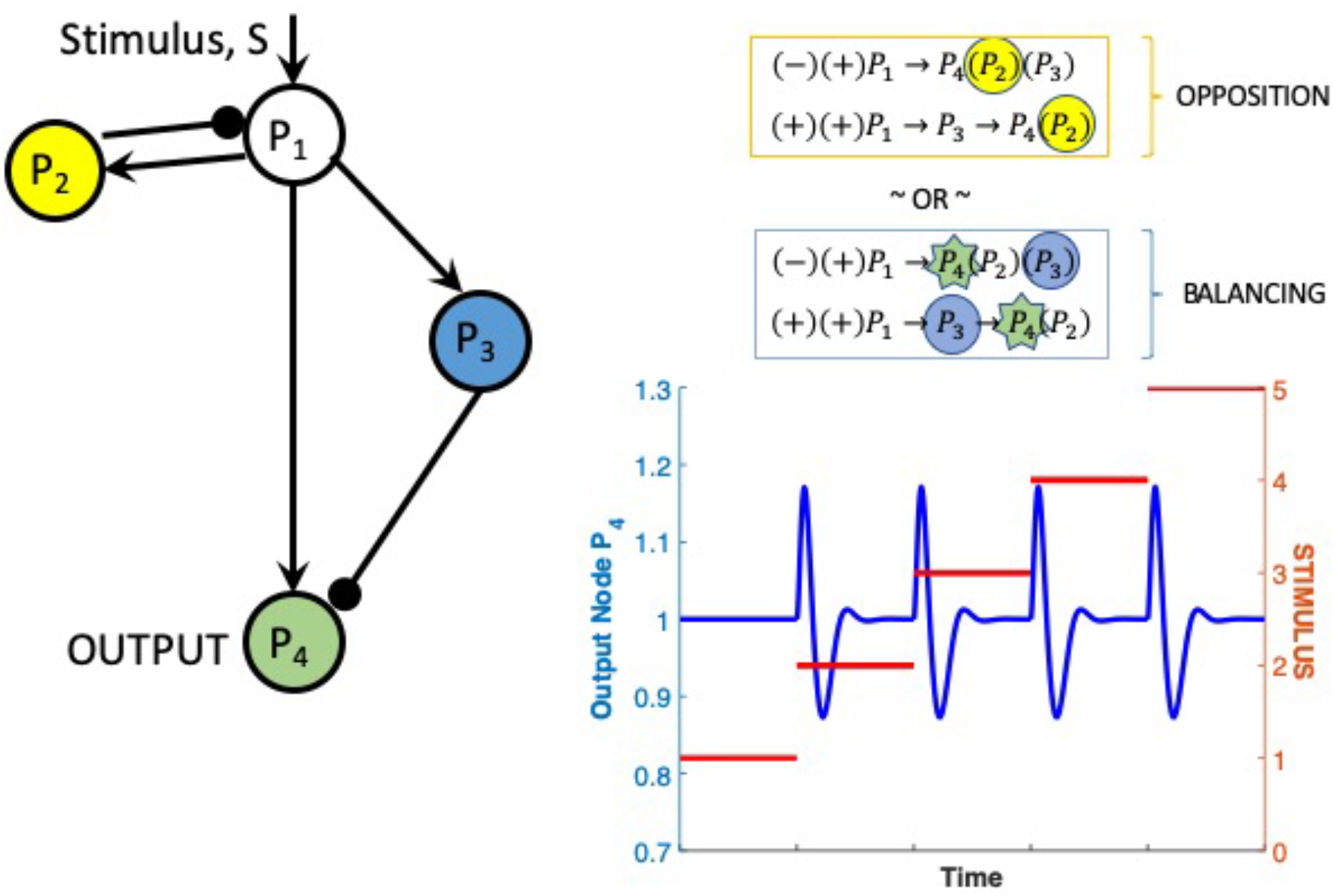
Network Example 2 – a two-node Opposer Module in series with a three-node Balancer Module. Parameters: k_1_ = k_2_ = k_3_ = k_4_ = k_5_ = k_6_ = k_7_ = k_8_ = k_9_ = 1.

This series connection of the two modules becomes clear when we examine the two-term RPA equation for this network (see diagrammatic representation in Figure 9).

We simulate a set of model equations for this network, corresponding to the configuration as a single Opposer module (with diagrammatic solution highlighted in yellow as an opposition mechanism). The only node subject to a special reaction constraint in this scenario is P_2_, for which we again choose the simple zero-order form to confer opposer kinetics:

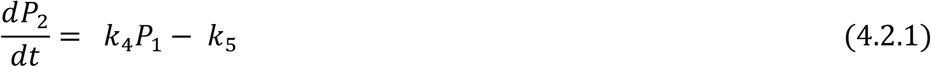

For the remaining three nodes, any reaction mechanism is allowed. We choose the following very simple rate equations:

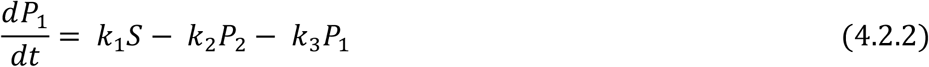

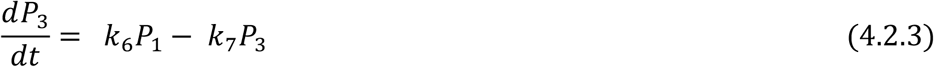

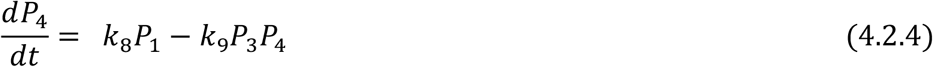

We note that the rate equations for P_3_ and P_4_ above happen to be consistent with balancer kinetics and connector kinetics, respectively. But any other functional forms for these equations would also be suitable, since the Opposer module is active as long as P_4_ operates with opposer kinetics, as reflected in Equation (4.2.1).

Simulation outputs for Equations (4.2.1) through (4.2.4) above are given in Figure 9. Note that for this particular choice of parameters (see Figure caption), the system exhibits damped oscillations. Indeed, as the stimulus undergoes a step increase, the output first increases rapidly, but then decreases and undershoots the setpoint. The signal then increases again and slightly overshoots the setpoint, before finally approaching the setpoint asymptotically. Since the set-point is independent of the stimulus magnitude, the system exhibits RPA. The time-dependent response of the system, on the other hand, and its propensity for oscillations, depends on the choice of parameters. Notice also that the time-dependent response exhibits a great deal of symmetry: as long as the difference in stimulus magnitude is preserved from one step-increase to the next, the time-dependent response of the system remains identical. This latter property is a consequence of the special (simple) choices of reaction kinetics used in this example.

### 4.3 Example Network 3: A single opposer module with two-node opposing set, and a single input-output node

In the simple four-node network example in Figure 10, we illustrate two interesting features that can characterise RPA-capable networks: (1) The network need not have distinct input-output nodes – a valid topological possibility rarely considered in the RPA literature to date; and (2) a single Opposer module may contain a whole constellation of opposer nodes, connected together in an orchestrated collection of interconnected feedback loops. For simplicity of illustration, we choose to examine a set of just two opposer nodes collaborating to confer RPA on the output node, P_1_ (which is also the node receiving the external stimulus, S).

**Figure 10:**
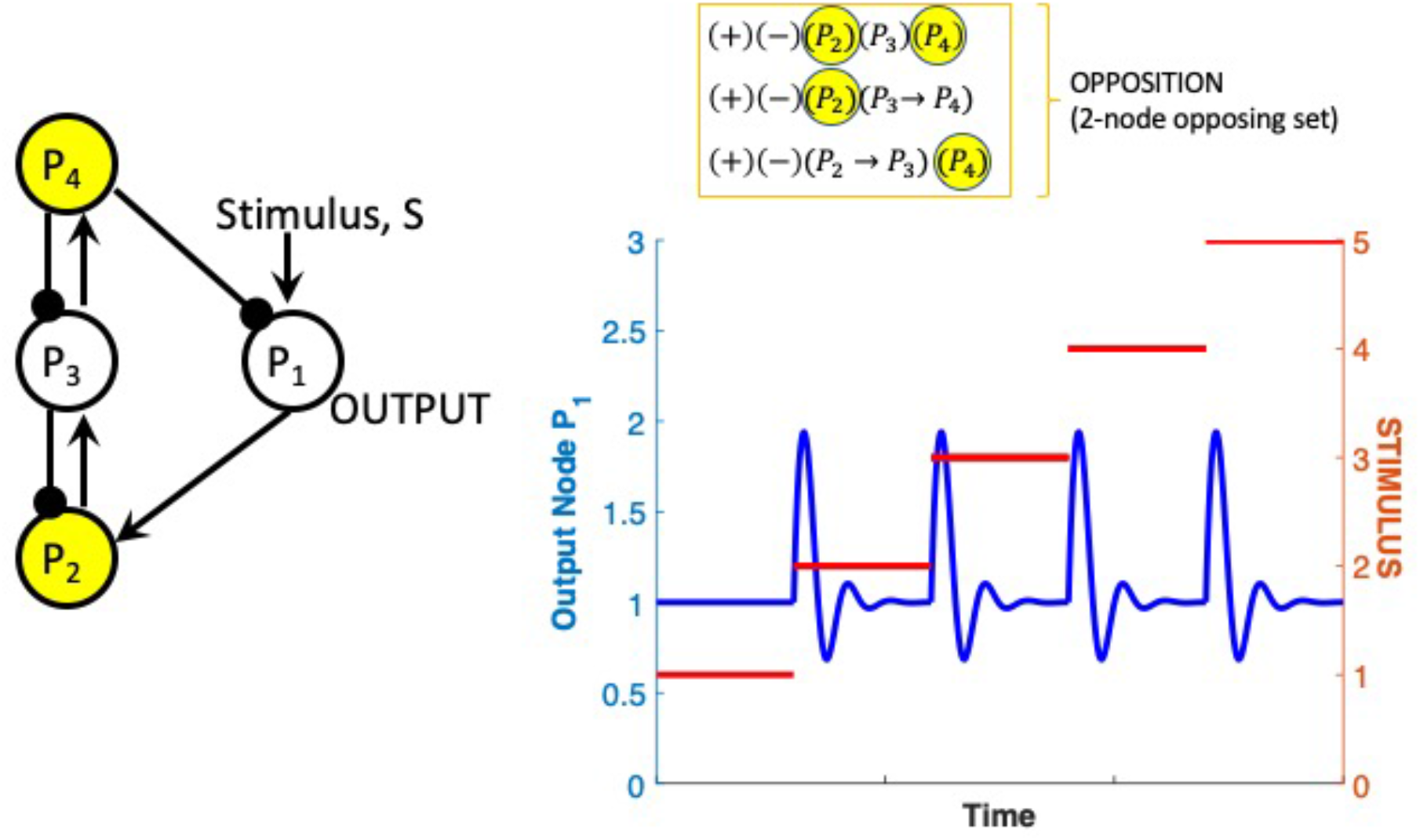
Network Example 3 – a two-node Opposing Set (a single Opposer Module with two collaborating opposer nodes). The network has a single input/output node (P_1_). Parameters: k_1_ = k_2_ = k_6_ = k_7_ = k_8_ = 1; k_3_ = k_4_ = k_5_ = k_9_ = k_10_ = 0.8.

Networks with a single input/output node have an RPA equation with a slightly different structure insofar as each term in the equation contains only the cycle component. There are no ‘routes’ in networks with a single input/output node. Thus, as noted in the summary of RPA network design features in Section 3.2, a network of this type can only achieve RPA through an Opposer Module. There are two (negative) feedback loops disjoint from the input/output node in this network, giving rise to three different cycle combinations as indicated in the diagrammatic representation in Figure 10. All three of the remaining nodes exist in feedback loops, and it is clear from the diagrammatic representation that the Opposer mechanisms 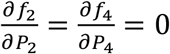 are sufficient to satisfy the RPA equation for this network.

We solve the following simple rate equations for this RPA network solutions for a range of increasing steps in the stimulus, S:

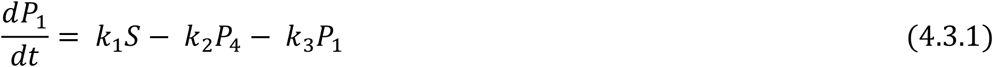

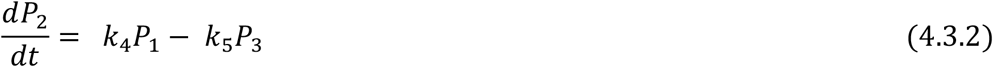

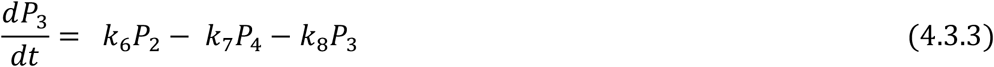

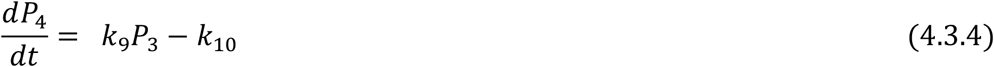

These solutions are depicted in Figure 10. Notice that for these simple reaction kinetics, and for the chosen parameters (see Figure caption), the time-dependent response exhibits damped oscillations as well as a high level of symmetry for fixed step-increases in stimuli.

### 4.4. Example Network 4: A single RPA-capable network with two possible RPA solutions

In the example depicted in Figure 11, we highlight two crucial properties that characterise many RPA-capable networks:

**Figure 11:**
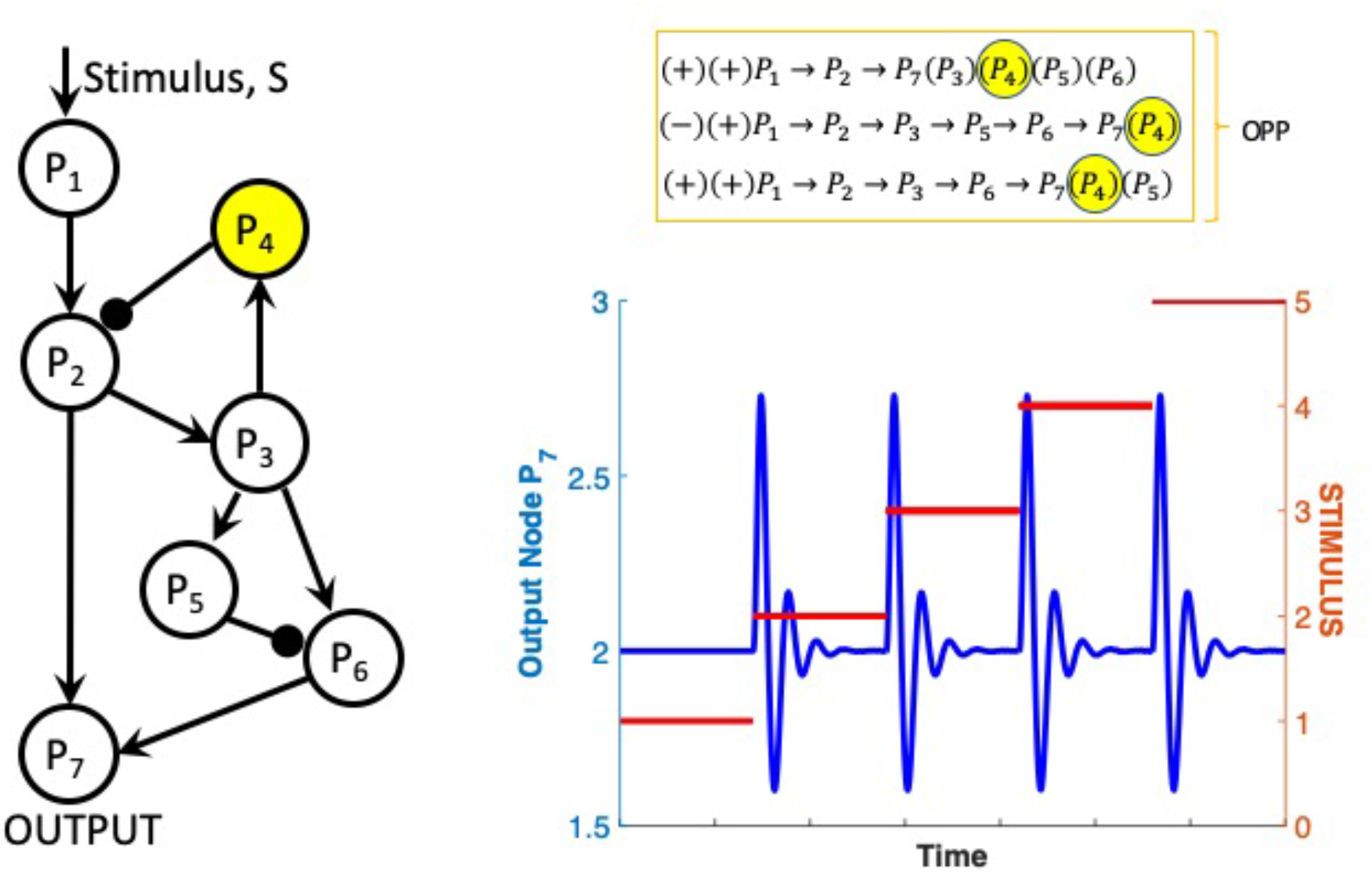
Network Example 4 – An RPA-capable network with two possible decompositions into basis modules. Here we depict the network in its realization as a single Opposer Module. Parameters: k_1_ = k_2_ = k_3_ = k_4_ = k_5_ = k_10_ = k_11_ = k_12_ = k_13_ = k_14_ = k_15_ = k_16_ = 1. k_6_ = k_7_ = k_8_ = k_9_ = 0.5.

1. RPA may be achieved in large networks with constraints at only a single node of the network. In other words, all but one of the vast array of chemical reactions comprising a complex network may be perturbed (mutationally, pharmacologically, or otherwise) while exerting no influence whatsoever on that network’s ability to exhibit RPA. Only the single computational node of the network (e.g. the opposer node P_1_ in the case of the network in Figure 11) is subject to a strict constraint on its reaction mechanism (here, an opposer mechanism).
2. For networks of sufficient complexity, there may be multiple ways for the network to be decomposed into RPA modules.

Although the network in Figure 11 contains seven interacting nodes its very simple architecture, and sparse input-output minor, endows it with a small RPA equation: Three distinct routes from P_1_ to P_7_ exist in this network (P_1_→P_2_→P_7,_ P_1_→P_2_→P_3_→P_6_→P_7_ and P_1_→P_2_→P_3_→P_5_→P_6_→P_7_), with the single network feedback loop (P_2_→P_3_→P_4_) contiguous with all three. The RPA equation is limited to the three terms noted in Figure 11. Since each term contains the kinetic multiplier (P_4_) in its cycle component, and the node P_4_ occurs in a feedback loop, this entire network can achieve RPA through the activity of the single opposer node P_4_ alone.

For illustration, we simulate a set of model equations for this RPA possibility, again choosing zero-order kinetics for the reaction rate at P_4_ to guarantee opposer kinetics for this node:

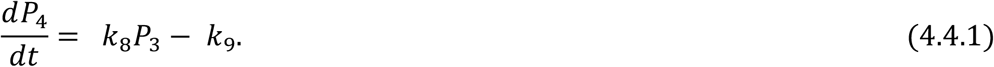

Once again, an arbitrary choice can be made for the remaining reaction rates, so we choose the following very simple forms:

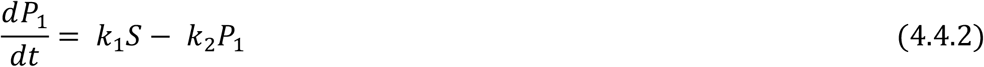

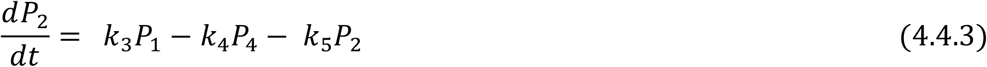

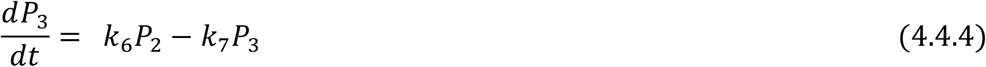

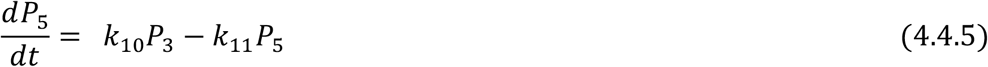

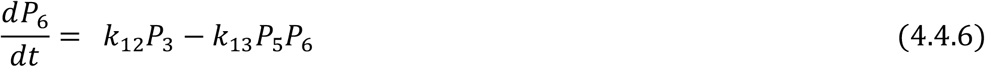

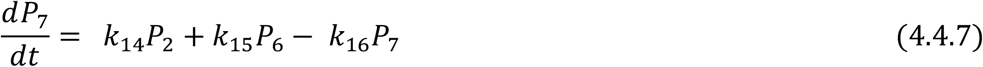

Solutions to these equations are given in Figure 11 (with parameter choices listed in the Figure caption). Here again, for the simple reaction kinetics we’ve employed, and for this particular parameter set, the time-dependent response exhibits damped oscillations as well as a high level of symmetry for fixed step-increases in stimuli.

Now, the node P_3_ also occurs in a feedback loop in this network, and is thus a candidate Opposer node. Nevertheless, its kinetic multiplier only appears in one of the three terms owing to the fact that P_3_ also occurs in a route of this network (as reflected in the remaining two terms). With P_3_ acting as an Opposer node, the network may still achieve RPA through the simultaneous activity of a Balancer module, with P_5_ as a Balancer node, and P_6_ as a Connector node. This gives rise to the two-module structure depicted in Figure 12, in which an Opposer module and a balancer module coexist in parallel.

**Figure 12:**
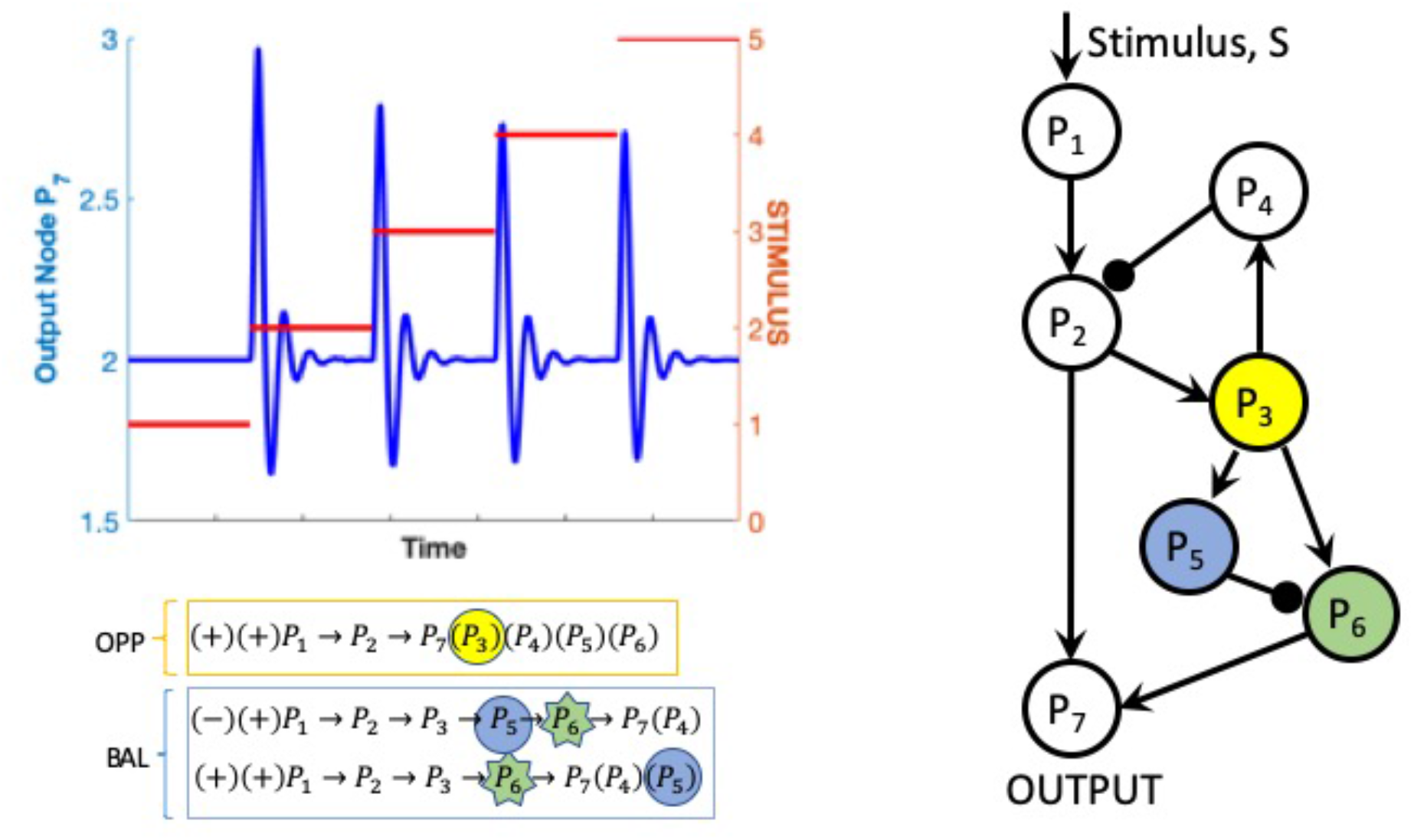
Network Example 4 with an alternative RPA solution. Here the network is shown as a three-node Opposer Module in parallel with a three-node Balancer Module. Parameters: k_1_ = k_2_ = k_3_ = k_4_ = k_5_ = k_10_ = k_11_ = k_12_ = k_13_ = k_14_ = k_15_ = k_16_ = 1. k_6_ = k_7_ = k_8_ = k_9_ = 0.5.

For this alternative RPA solution, with P_3_ acting as an Opposer node, the node P_4_ may no longer possess opposer kinetics. We thus replace the rate equation for this node with the following simple form:

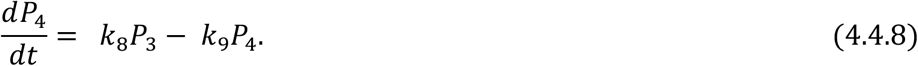

We also replace the rate equation for P_3_ with a zero-order rate law for which opposer kinetics are assured:

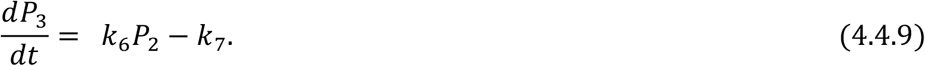

Moreover for this two-module RPA solution, P_5_ must operate with balancer kinetics, and P_6_ with connector kinetics. We observe that Equations (4.4.5) and (4.4.6) already conform to these requisite reaction mechanisms (although no special reaction kinetics were required at these nodes for the previous RPA solution noted above). We solve the new set of model equations (incorporating the new rate equations (4.4.8) and (4.4.9) above) and display the results in Figure 12. Notice that the symmetry that previously characterized the time-dependent solutions (Figure 11) has now been lost in this two-module version of the network due to the alterations in reaction mechanisms.

### 4.5 Example Network 5: A twelve-node RPA-capable network with two opposer modules in parallel

In our final example, we emphasize again that RPA may achieved in larger networks with only very few constraints on individual nodes. By the diagrammatic method we’ve demonstrated throughout

this chapter, it is straightforward to show that the simple twelve-node network configuration illustrated in Figure 13 has an RPA equation with fourteen terms. As we highlight in the diagrammatic representation of this RPA equation, the kinetic multiplier (P_9_) occurs in eight of these terms, while the kinetic multipliers (P_5_) and (P_6_) occur in the remaining six terms. Nodes P_5,_ P_6_ and P_9_ are all contained in feedback loops, so they are suitable choices for Opposer nodes. By proposing a set of simple modeling equations that endow, with P_9_ subjected to an opposer reaction mechanism, and either P_5_ or P_6_ also subjected to an opposer mechanism, we show in Figure 13 that this network readily exhibits RPA through this decomposition into two parallel Opposer modules.

**Figure 13:**
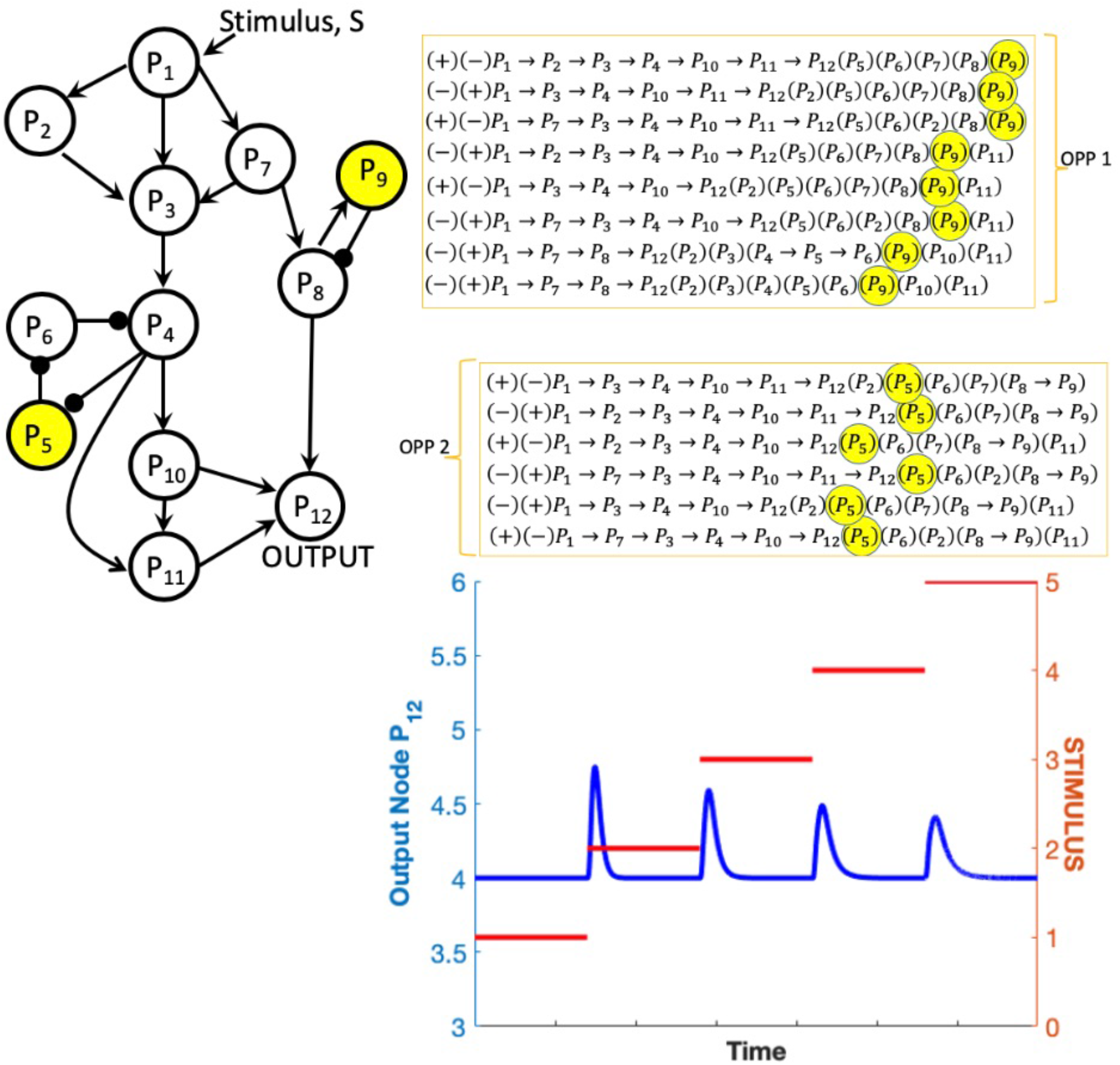
Network Example 5 – A twelve-node RPA-capable network with two Opposer Modules in parallel. Parameters: k_1_ = k_2_ = k_3_ = k_4_ = k_5_ = k_6_ = k_7_ = k_8_ = k_15_ = k_16_ = k_22_ = k_23_ = k_24_ = k_25_ = k_26_ = k_27_ = k_28_ = k_29_ = k_30_ = 1; k_9_ = k_10_ = k_11_ = k_12_ = k_13_ = k_14_ = k_17_ = k_18_ = k_19_ = k_20_ = k_21_ = 1.

We choose the following simple set of equations for the model simulations in Figure 13:

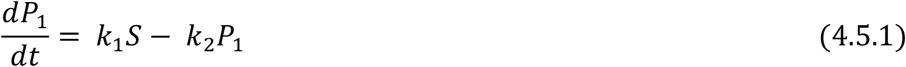

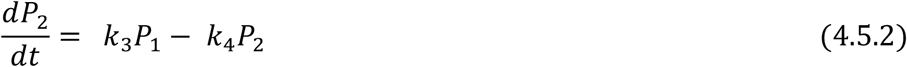

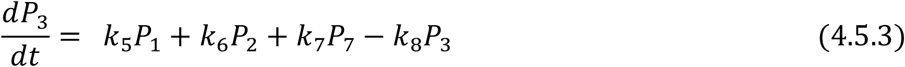

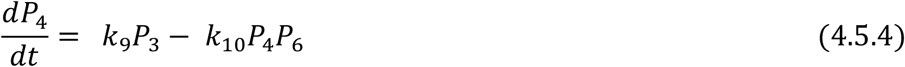

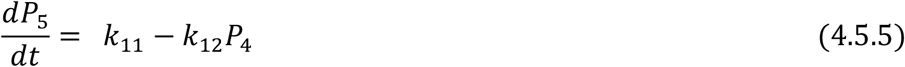

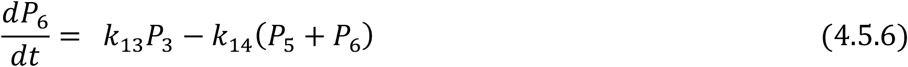

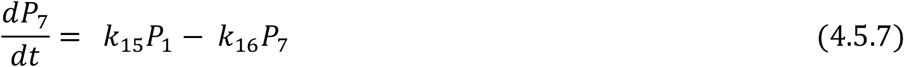

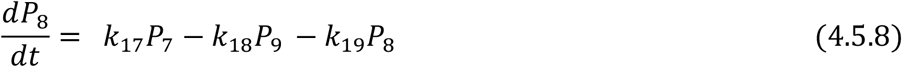

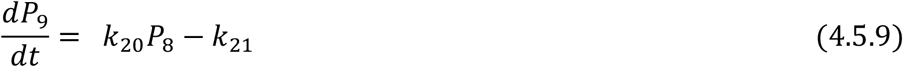

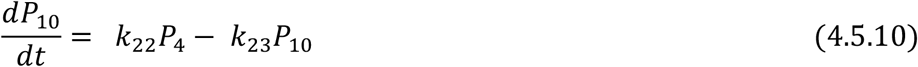

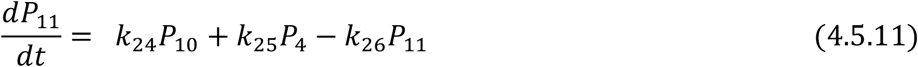

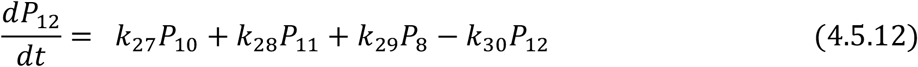

Notice that for this particular choice of parameters, no oscillations are observed in the output signal in response to each step-increase in stimulus. Indeed, the output decreases monotonically to the fixed stimulus-independent baseline level after its initial transient increase.

## 5. Concluding Remarks: Future Directions for RPA Theory

The analysis of the complex networks arising in nature confronts us with extraordinary technical challenges. In many biological contexts – cellular signal transduction and cellular metabolism, for instance – the underlying signaling networks are so complex and high-dimensional that their detailed topologies and reaction mechanisms are essentially unknowable. Without access to the full network design space for a particular functional requirement, proposing an accurate and predictive mathematical model for a particular network, or testing such a network to see if it meets a particular mathematical criterion, would require us to know (a) *all the molecules* involved in the cascades of chemical reactions that are initiated once a “stimulus” of some kind is delivered to the system (including all scaffolds, adaptor proteins, second messengers and other ancillary molecules), and (b) the reaction mechanisms (e.g. rate laws and kinetics) of all reactions, including the role of intermediate complexes in all enzyme-catalysed reactions, and the specific mechanisms of multisite covalent modification (which could be processive or distributive in nature, for example). In practice, we would also require information on the abundances of all reactant molecules, since the kinetics of a given reaction can vary in important qualitative ways as the involved enzymes become “saturated” by their substrates.

The topological method we discuss here for solving the RPA problem is thus an important step forward in our quest to determine how biological complexity is organised. Indeed, the strict mathematical requirement for robust networks to exist in well-defined modular structures suggests the tantalising concept that complex networks are like snowflakes: while each is unique, all individual instances are alike in their essential structure. Our hope is that the pragmatic discussion of the general RPA solution presented here will stimulate broad interest in these new approaches, and promote the discovery of new robustness-preserving network designs for other functional requirements of biological systems.

Before closing, we wish to highlight a frontier research problem in RPA theory: the deep question of how integral control is implemented at the microscale of complex networks through intricate intermolecular interactions. Although we have employed very simple reaction rate laws for illustrative purposes in our example networks here, the molecular implementation of integral control in intracellular compartments is almost certainly vastly more complex. Significant progress has been made in this direction through the study of a special case of RPA known as absolute concentration robustness (ACR). It is now clear *[5]* that the Shinar-Feinberg model of ACR *[30]* is a simple balancer module, although the connector kinetics required for this model are buried much more deeply into the reactions of this model than the simple connector mechanisms employed in this chapter. Even the Opposer node in the antithetical integral control model *[22]* requires a transformation of coordinates to achieve opposer kinetics *[31]*. In addition, the deep connection between RPA and ultrasensitivity [5] suggests a wealth of future avenues for identifying new mechanisms for the construction of robustness-promoting integrals from amid intricate biochemical reactions. The study of robustness at network microscales along these lines will offer us fresh insights into the biological signalling mechanisms underpinning human disease states, along with new possibilities for pharmacological disruption, and will spawn a host of new opportunities in synthetic biology.

## Acknowledgements

Robyn P. Araujo is the recipient of an Australian Research Council (ARC) Future Fellowship (project number FT190100645) funded by the Australian Government.

